# Prevalent, protective, and convergent IgG recognition of SARS-CoV-2 non-RBD spike epitopes in COVID-19 convalescent plasma

**DOI:** 10.1101/2020.12.20.423708

**Authors:** William N. Voss, Yixuan J. Hou, Nicole V. Johnson, Jin Eyun Kim, George Delidakis, Andrew P. Horton, Foteini Bartzoka, Chelsea J. Paresi, Yuri Tanno, Shawn A. Abbasi, Whitney Pickens, Katia George, Daniel R. Boutz, Dalton M. Towers, Jonathan R. McDaniel, Daniel Billick, Jule Goike, Lori Rowe, Dhwani Batra, Jan Pohl, Justin Lee, Shivaprakash Gangappa, Suryaprakash Sambhara, Michelle Gadush, Nianshuang Wang, Maria D. Person, Brent L. Iverson, Jimmy D. Gollihar, John Dye, Andrew Herbert, Ralph S. Baric, Jason S. McLellan, George Georgiou, Jason J. Lavinder, Gregory C. Ippolito

## Abstract

Although humoral immunity is essential for control of SARS-CoV-2, the molecular composition, binding epitopes and effector functions of the immunoglobulin G (IgG) antibodies that circulate in blood plasma following infection are unknown. Proteomic deconvolution of the circulating IgG repertoire (Ig-Seq^1^) to the spike ectodomain (S-ECD^2^) in four convalescent study subjects revealed that the plasma response is oligoclonal and directed predominantly (>80%) to S-ECD epitopes that lie outside the receptor binding domain (RBD). When comparing antibodies directed to either the RBD, the N-terminal domain (NTD) or the S2 subunit (S2) in one subject, just four IgG lineages (1 anti-S2, 2 anti-NTD and 1 anti-RBD) accounted for 93.5% of the repertoire. Although the anti-RBD and one of the anti-NTD antibodies were equally potently neutralizing *in vitro*, we nonetheless found that the anti-NTD antibody was sufficient for protection to lethal viral challenge, either alone or in combination as a cocktail where it dominated the effect of the other plasma antibodies. We identified *in vivo* protective plasma anti-NTD antibodies in 3/4 subjects analyzed and discovered a shared class of antibodies targeting the NTD that utilize unmutated or near-germline IGHV1-24, the most electronegative IGHV gene in the human genome. Structural analysis revealed that binding to NTD is dominated by interactions with the heavy chain, accounting for 89% of the entire interfacial area, with germline residues uniquely encoded by IGHV1-24 contributing 20% (149 Å^2^). Together with recent reports of germline IGHV1-24 antibodies isolated by B-cell cloning^3,4^ our data reveal a class of shared IgG antibodies that are readily observed in convalescent plasma and underscore the role of NTD-directed antibodies in protection against SARS-CoV-2 infection.

## INTRODUCTION

SARS-CoV-2, the causative agent of the COVID-19 pandemic, has spread to every continent except for Antarctica. The spike (S) surface glycoprotein is the primary antigenic target for the majority of vaccines and monoclonal antibodies (mAbs) currently under development or in clinical trials worldwide^5-8^. The S ectodomain (S-ECD) folds into a multidomain architecture^2,9^ and includes the angiotensin-converting enzyme 2 (ACE2) receptor binding domain (RBD), which is essential for viral infectivity, and the structurally adjacent N-terminal domain (NTD), which plays an uncertain role.

MAbs targeting the spike have been isolated predominantly by single B-cell cloning followed by screening for binding and neutralization *in vitro*^*10-18*^. These studies have revealed B-cell recognition of multiple spike epitopes and have led to the discovery of neutralizing antibodies primarily targeting the RBD. By contrast, the epitopes targeted by the circulating antibodies in convalescent individuals, especially those IgG plasma antibodies that are most abundant and that play a dominant role in the protective humoral immune response, have not been reported. Whereas serological assays of COVID-19 plasma have demonstrated that both the IgG plasma antibody repertoire and the B-cell repertoire can recognize multiple spike epitopes^19-22^, the fact that the clonal diversity and temporal dynamics of the two repertoires are divergent is not widely appreciated in general and very little is known about this important facet of antibody immunity in COVID-19^1,23-25^. For example, evidence for divergent repertoires in COVID-19 convalescents includes the paradoxical observation that B cells specific for S or RBD may be detected at high frequency in a given individual from whom potent virus-neutralizing mAbs can be cloned, yet their plasma virus-neutralizing activity remains low or negligible^10,21,26,27^. Here we report on the detailed and quantitative analysis of the molecular composition, epitope specificity and contribution to protection of the constituent antibodies that comprise the serological repertoire in COVID-19 convalescent study subjects.

## RESULTS

### Cohort & methodology

Blood was collected between days 11–19 post-onset of symptoms from four seroconverted convalescent study subjects with PCR-confirmed SARS-CoV-2 infections who experienced mild to moderate COVID-19 disease. Plasma virus-neutralizing titers (ID_50_s) with authentic SARS-CoV-2 USA-WA1/2020 ranged from <1:10 to approximately 1:285 (**Extended Data Fig. 1**). The lineage composition and relative abundance of IgG antibodies comprising the plasma response to either intact stabilized S-ECD (S-2P^2^) or RBD was determined using the Ig-Seq pipeline^1,23,28,29^ that integrates LC–MS/MS proteomics of affinity chromatography–enriched IgG antibodies with peripheral B-cell heavy-chain (V_H_), light-chain (V_L_), and single B-cell V_H_:V_L_ variable region repertoires (BCR-Seq) (**Fig. 1a)**.

**Figure 1.**
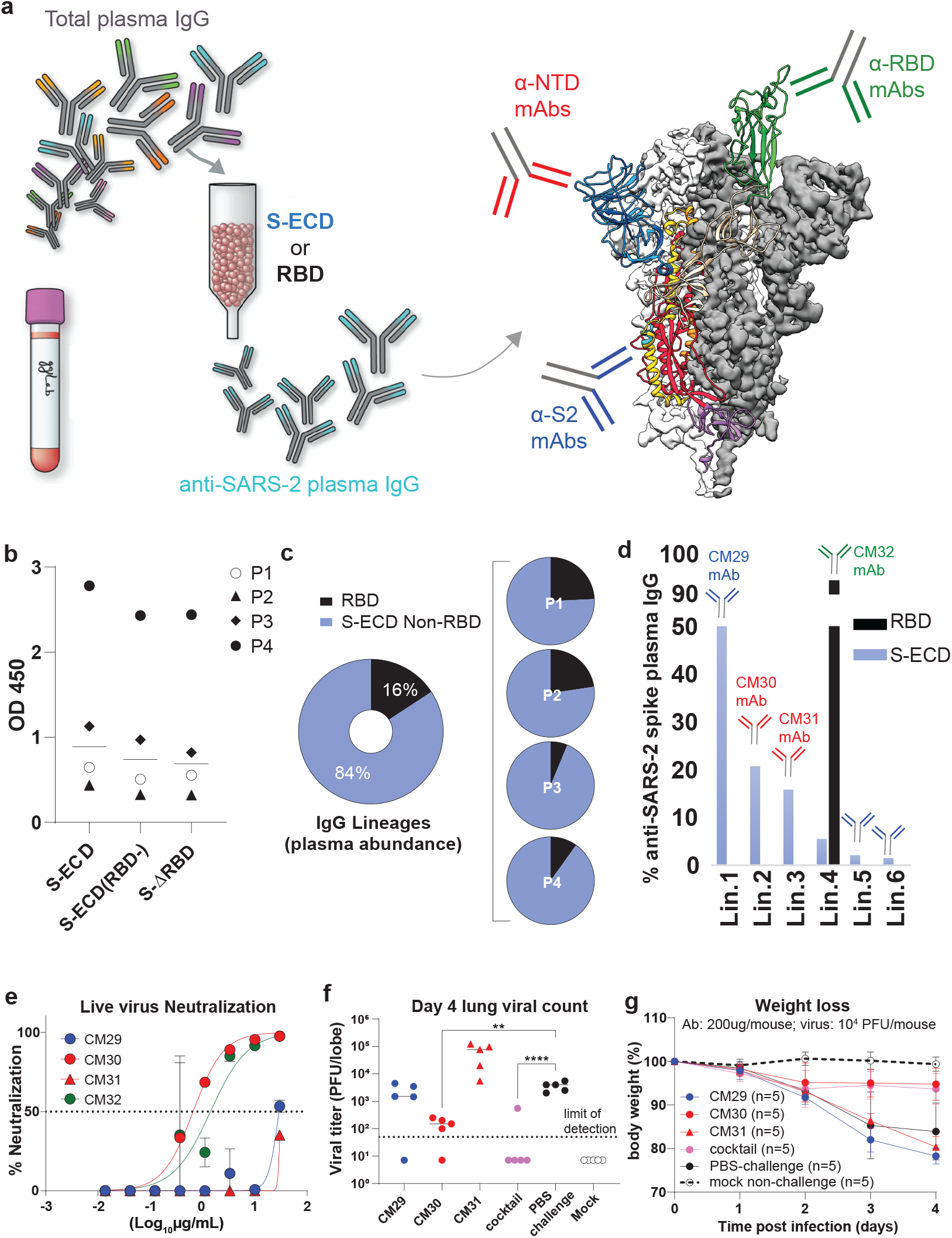
A single spike NTD-targeting IgG antibody in plasma can confer protection without a need for RBD-directed activity. **a**, Polyclonal IgG plasma antibodies were affinity purified using stabilized spike S-2P S-ECD^2^ or RBD, and binding specificity was mapped using purified S subdomains; anti-RBD (green); anti-S2 (blue); anti-NTD (red). **b**,The majority of the plasma anti-S-ECD response is directed to non-RBD epitopes: Binding (1:150 plasma dilution) to S-ECD alone, or in the presence of 50 µg/mL (∼1.7 µM) RBD, or S-ΔRBD deletion mutant. **c**, Quantitative determination of plasma anti-RBD and non-RBD antibody abundance in early convalescence. Abundance normalized to the entire anti-S-ECD plasma IgG repertoire. **d**, Composition, binding specificity and relative abundance of antibodies in early convalescent plasma (subject P3). **e**, Authentic virus neutralization of the four topmost abundant plasma IgGs (CM29, CM30, CM31, CM32) from plasma lineages Lin.1, Lin.2, Lin.3, Lin.4 in **1d** that account for >90% of the plasma anti-S-ECD repertoire. **f**,**g** Prophylactic protection of 12 m.o. BALB/C mice against lethal challenge with 10^4^ PFU mouse-adapted (MA10) SARS-CoV-2 using 200µg/mouse of non-RBD mAbs CM29, CM30, and CM31. Antibody cocktail (200 µg/mouse) consisted of 2:1:1 ratio of CM29, CM30, CM31, reflecting their relative plasma abundance (**1d**). \*\**p*<0.005; \*\*\*\**p*<0.0001.

### Plasma IgG repertoire composition and function

To analyze the composition of the polyclonal IgG response directed against the S glycoprotein in convalescent plasma, we performed differential pulldowns to fractionate S-ECD-directed versus RBD-directed antibody repertoires (**Fig. 1a**). Antibody lineages detected by Ig-Seq in the S-ECD affinity chromatography eluant but absent from the RBD eluant were deemed to be reactive with S epitopes outside the RBD. In ELISA assays plasma IgG binding to S-ECD and to S-ECD ΔRBD deletion mutant was statistically similar (p>0.05), and a minimal drop in binding signal to S-ECD was also observed in the saturating presence of ∼1.7 µM purified RBD as competitor (**Fig. 1b**). Molecular-level serology using Ig-Seq confirmed this observation (**Fig. 1c; Extended Data Fig. 2**) and further revealed across the cohort that, on average, 84% of the response to S-ECD bound to epitopes outside the RBD. In all subjects, the detected plasma IgG repertoire to S-ECD was oligoclonal comprising between only 6–22 lineages, with the top-ranked lineage comprising 15%-50% abundance of the entire repertoire (integrated LC-MS/MS XIC intensity). Earlier studies have estimated that the Ig-Seq pipeline typically captures >70% of the circulating virus-specific antibody repertoire, which can increase to >85% identification for the most abundant lineages^30^. Furthermore, in all four subjects the topmost abundant plasma IgG lineage was directed to epitopes in S-ECD outside the RBD (**Fig. 1d, Extended Data Fig. 2**). These patterns were uniformly observed across all the study subjects irrespective of differences in virus-neutralizing titer (Extended Data Fig. 1) or S-binding titer (**Fig. 1b**). This pattern of serological reactivity agrees with a prior study of B-cell responses and the cloning of 2045 mAbs, most of which bound to regions of the spike outside of the RBD, indicating that viral infection generates a strong response against non-RBD epitopes^15^.

In subject P3, whose disease resolved within two days post-onset of symptoms and who had the lowest plasma neutralizing titers (**Extended Data Fig. 1**), we detected six IgG lineages binding to S-ECD (**Fig. 1d**). Furthermore, only four of the lineages (Lin.1–Lin.4) were detectable at levels >5% abundance of the total S-ECD repertoire. Of these, the top two IgG lineages when combined accounted for >70% of the total S-ECD response. For each of these top two plasma IgG lineages (Lin.1 and Lin.2), we detected abundant intra-lineage diversity with 22 and 11 unique complementarity-determining region H3 (CDR-H3) peptides mapping to each lineage, respectively (**Extended Data Fig. 3**) indicating somatic hypermutation and clonal expansion. In total, Lin.1–Lin.4 accounted for 93.5% abundance by integrated XIC intensity suggesting that they overwhelmingly account for the detectable S-ECD plasma repertoire.

ELISA and biolayer interferometry analysis of mAbs CM29–CM32 representing the most expanded clones within each of Lin.1–Lin.4 lineages showed that CM29 (Lin.1) recognizes the S2 subunit (K_D_ = 6.6 nM), CM30 and CM31 (Lin.2 and Lin.3 with K_D_ = 0.8 and 37.7 nM, respectively) were specific for the NTD, and CM32 (Lin.4 K_D_ = 6.0 nM) bound the RBD, as expected from the Ig-Seq differential affinity-chromatography pulldowns (**Fig. 1d, Extended Data Table 1**). The anti-NTD antibody CM30 potently neutralized authentic SARS-CoV-2 (IC_50_ 0.83 μg ml^-1^), the anti-RBD CM32 was slightly less potent (2.1 μg ml^-1^), and CM29 and CM31 showed very low or no neutralization activity (**Fig. 1e**).

We then determined the capacity of mAbs CM29–CM32, singly and in combination, to confer prophylactic protection to virus challenge using the MA10 mouse model^31,32^. This SARS-CoV-2 model exhibits the spectrum of morbidity and mortality of COVID-19 disease as well as aspects of host age, cellular tropisms, and loss of pulmonary function. Even though the RBD-directed mAb CM32 was potently effective for live-virus neutralization *in vitro* and had high antibody-dependent cellular phagocytosis (ADCP) activity relative to the other plasma mAbs from this subject (**Extended Data Fig. 4**) it did not confer protection *in vivo* in mice challenged with a low-dose infection (10^3^ plaque forming units (PFU)) nor did it reduce lung titers (**Extended Data Fig. 5**). Similarly, no protection was observed with S2-directed, non-neutralizing CM29 although a statistically significant −1.5 log_10_ reduction in viral lung titers was observed (*p*<0.0001), an effect that may be related to its moderately high ADCP activity (**Extended Data Fig. 4**). CM30, derived from the top-ranking NTD-targeting IgG lineage (21% abundance), was the sole plasma mAb (among those lineages detected at >5% of the S-ECD response) that conferred complete protection both in low MA10 viral load (**Extended Data Fig. 5**) and, most notably, with high viral load challenge (10^5^ PFU/mouse) (**Figs. 1f,g**). Interestingly, administration of the non-neutralizing anti-NTD plasma mAb CM31 conferred no protection and resulted in increased lung titers, although not statistically significant (mean increase: 11-fold, *p*=.08). Administration of a cocktail comprising the top non-RBD plasma mAbs CM29–CM31, which collectively represented >85% of the IgG plasma lineages to S-ECD (**Fig. 1b**), showed complete protection and lung viral titers below LOD in high viral load challenge. Combined, the protection data suggest that the presence of a single potent plasma non-RBD mAb is sufficient to counter the effect of other subprotective IgG antibodies in polyclonal plasma.

### Prevalent IgG antibodies target non-RBD epitopes

Subject P2, whose illness was more severe and protracted compared to subject P3, displayed a more polyclonal IgG response (**Fig. 2a**) yet 12/15 lineages comprising the anti-S-ECD repertoire bound to non-RBD S-ECD epitopes and accounted for >80% of the repertoire. This estimate was validated by analysis of the fine specificity of individually cloned recombinant plasma mAbs for the 8 of these 15 lineages for which we had paired VH:VL mAb sequence data, including all those detected at >5% abundance (collectively accounting for >75% of the response) (**Fig. 2a**). Interestingly, as with P3, the most abundant S-ECD plasma antibodies target the S2 subunit, with the four topmost lineages (68% total abundance) binding to S2. Broadly reactive functional antibody responses against S-ECD, but not RBD, have been noted to be enriched and trend with protection from both primary disease and reinfection in humans and non-human primates^33,34^. We noticed that mAbs CM25 and CM17, representative of two NTD-targeting lineages each comprising ∼2.5% of the response at day 56 (Ig-Seq Lin.6 and Lin.9; **Fig. 2a**), were both encoded by germline or near-germline IGHV1-24. We also found an additional NTD-targeting IGHV1-24 plasma mAb (CM58) in subject P4.

**Figure 2.**
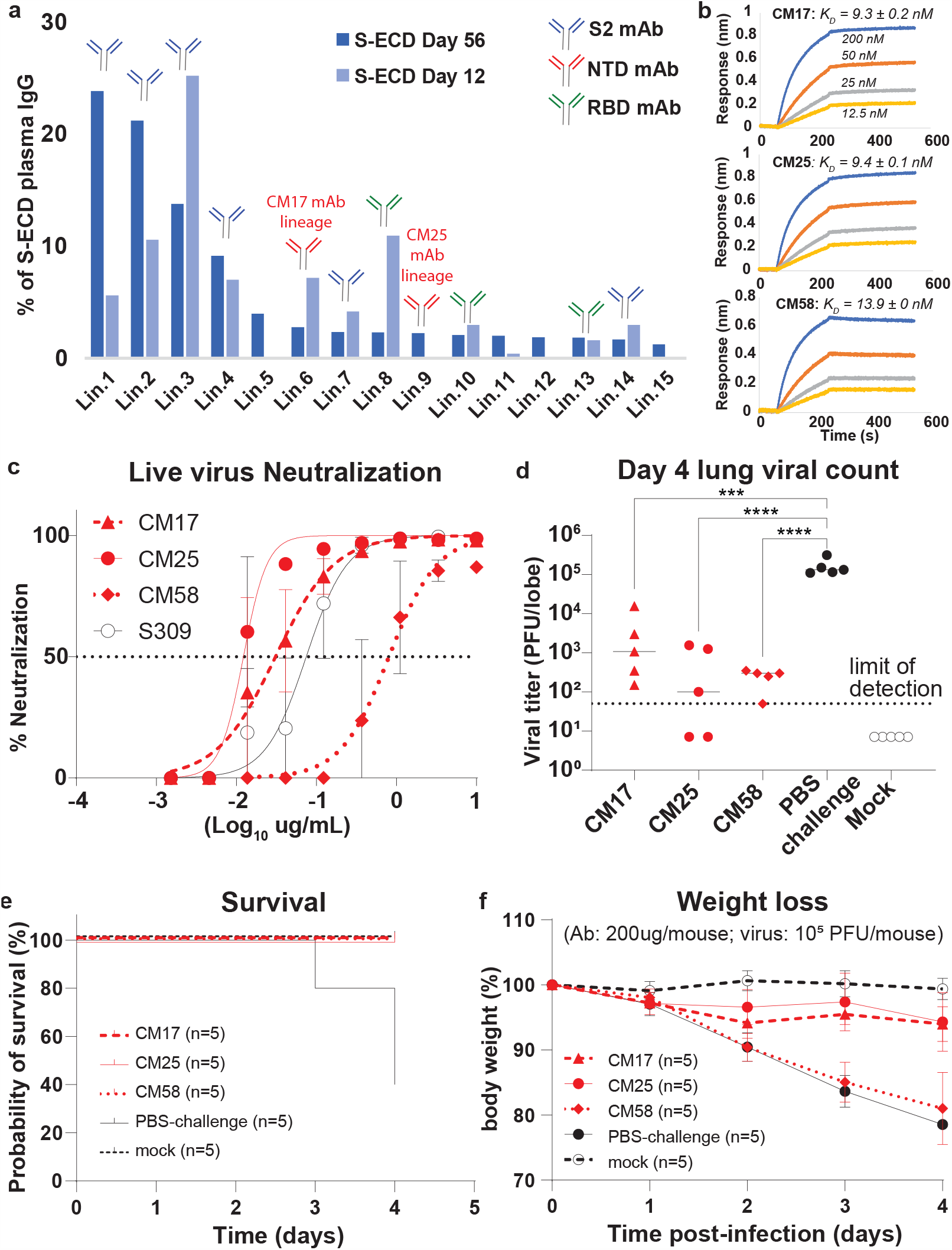
Protective spike NTD-targeting antibodies are prevalent in the plasma of convalescent COVID-19 study subjects. **a**, Temporal dynamics of the anti-S-ECD IgG repertoire at days 12 and 56 post-symptom onset. Titer had increased two-fold by day 56 (data not shown). **b**, Biolayer interferometry (BLI) sensorgrams to S-ECD ligand of anti-NTD mAbs CM17 and CM25 (subject P2), CM58 (subject P4), and anti-RBD control mAb S309^35^. **c**, *In vitro* live virus neutralization. **d–f**, Prophylactic protection of 12 m.o. BALB/C mice against intranasal challenge with 10^4^ PFU of mouse-adapted (MA10) SARS-CoV-2. *In vivo* prophylactic protection to MA10 challenge; experimental conditions as in **Fig 1f,g** except challenge dose was 10^5^ PFU. \*\*\**p*<0.0007; \*\*\*\**p*<0.0001.

CM17, CM25 and CM58 bound S-ECD with nM affinities (**Fig. 2b; Extended Data Table 1**). In two independent assays using two different Vero-E6 cell sublines, all three of these plasma IGHV1-24 mAbs potently neutralized SARS-CoV-2 virus, with IC_50_ 0.01, 0.03 and 0.81 μg ml^-1^ for CM25, CM17 and CM58 respectively (**Fig. 2c; Extended Data Fig. 6; Extended Data Table 1**). CM25 in particular has potency comparable to, or exceeding, clinical-stage antibodies targeting the RBD^13,16,35,36^ (**Fig. 2c**). For all three mAbs, pre-administration in the MA10 mouse model resulted in significantly reduced lung viral titers at day 4 post-infection with 10^5^ PFU of live virus (**Fig. 2d**; *p*<0.001). At day 4, all mice in the antibody treated groups survived, compared to just 40% in the control group (**Fig. 2e**); however, animals treated with CM58 experienced significant weight loss, while those treated with either CM25 or CM17 did not (**Fig. 2f**).

### Genetic convergence of IGHV1-24 NTD antibodies

While IGHV1-24 is expressed in the B-cell repertoire of healthy individuals at a relatively low frequency (0.4%–0.8%),^37^ this gene segment has been observed to be expressed ∼10-fold higher than expected (∼5%-8%) in memory B cell and plasmablast repertoires from COVID-19 patients^4,12,38^. This prompted us to examine whether IGHV1-24 frequency might similarly be elevated at the serological level in COVID-19 convalescent plasma. Examination across all four study subjects detected IGHV1-24 antibodies exclusively in the S-ECD affinity chromatography eluant (median 4.3%) when compared with the RBD eluant (0%) (**Fig. 3a,b**), indicating that IGHV1-24 partitions exclusively with IgG recognition of S-ECD epitopes that lie outside the RBD (**Fig. 3a**). Notably, IGHV1-24 antibodies likely recognizing unrelated antigens were detected in the non-binding flow-through chromatography fraction, albeit at lower frequency (**Fig. 3b**). Whereas most IGHV1-24 plasma antibodies detected in COVID-19 convalescent plasmas bind to S-ECD, we note that serological responses directed against the related class I viral membrane fusion (spike) proteins of influenza and respiratory syncytial virus are not enriched for IGHV1-24 **(Fig. 3b**).

**Figure 3.**
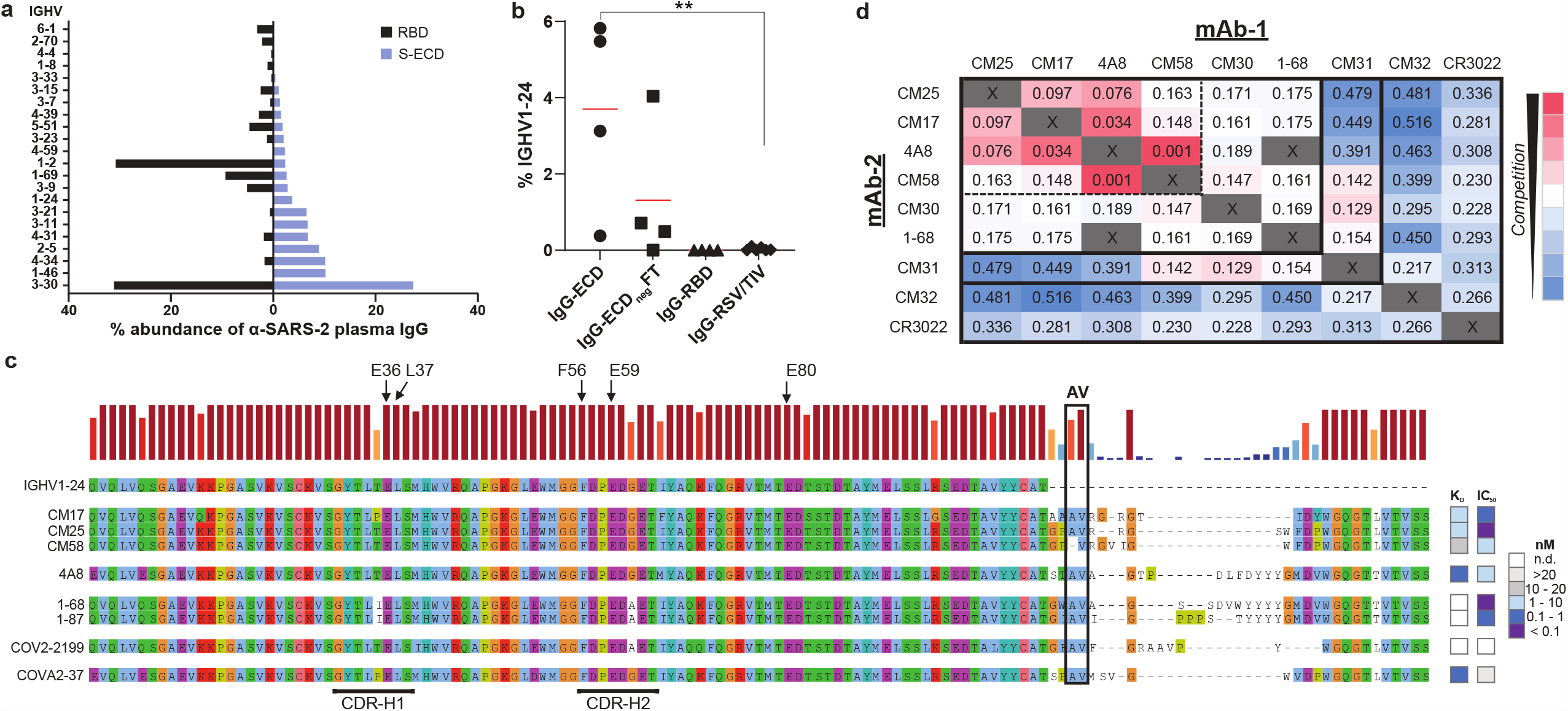
Genetic basis of a shared, or public, class of IGHV1-24 plasma antibodies targeting the spike NTD. **a**, IGHV usage of plasma anti-S-ECD or anti-RBD antibodies in all subjects (n=4). **b**, IGHV1-24 antibodies as a percentage of the circulating IgG plasma antibody repertoire: reactivity to SARS-CoV-2 spike S-ECD or RBD in COVID-19 subjects, or reactivity in healthy subjects to vaccine spike antigens for either respiratory syncytial virus (RSV) or trivalent influenza vaccine hemagglutinin HA1 (TIV). \*\**p*<0.01. **c**, Sequence alignment of IGHV1-24 neutralizing anti-NTD IgGs from plasma (CM25, CM17 and CM58) or from peripheral B cells (4A8^3^, 1-68 and 1-87 from a subject with ARDS^4^, COV2-2199^13^, and COVA2-37 [mild disease subject])^12^. Arrows point to unique IGHV1-24 residues. Heatmap shows recombinant mAb affinity (K_D_) and live virus neutralization (IC_50_) for individual antibodies. **d**, Competitive BLI binding assay (“checkboard competition”) of NTD-binding mAbs found in this study (CM17, CM25, CM58, CM30, and CM31) and others (4A8 and 1-68). RBD-binding mAbs CM32 and CR3022 included for comparison. Numbers refer to the shift, in nanometers, after second mAb binding to the preformed mAb–NTD complex.

Multiple alignment (**Fig. 3c)** of the plasma mAbs CM17, CM25 and CM58 with four recently reported neutralizing IGHV1-24 anti-NTD mAbs cloned from peripheral B cells (4A8^3^, 1-68^4^, 1-87^4^, COVA2-37^12^) and an additional antibody (COV2-2199^13^) for which no neutralization data had been reported identified a pattern of highly similar V_H_ immune receptor sequences (**Fig. 3d**). The V_H_ region of all 8 mAbs exhibits zero or low somatic mutation (nucleotide identities 97– 100%). In all cases, three glutamate (Glu) residues (Glu36, Glu59, Glu80; IMGT numbering^39^) located in CDR-H1, CDR-H2 and framework H3 (FWR-H3), respectively, as well as a phenylalanine (Phe) residue (Phe56) in CDR-H2, were invariably unmutated; these amino acids at these precise positions are unique to IGHV1-24, which occurs as a single nonpolymorphic allele among the 129 IGHV genes and alleles residing in the human genome^39^. Glu36, Glu59, and Glu80 occur in fewer than 1%, 2%, and 1% of 87,838 human antibody heavy chains respectively, while Phe56 occurs in fewer than 1.2%^40^. CDR-H3 peptide intervals were restricted to lengths of 14 or 21 amino acids and contained a key dipeptide motif near the amino terminus: an aliphatic AV patch (Ala109 and Val110) (**Extended Data Table 2**). Despite these constraints on the heavy chain, six different light chain V_L_ genes are observed (**Extended Data Table 2**) suggesting a minor role for the V_L_ domain for NTD binding activity or specificity. A “checkerboard” binding-competition experiment (**Fig. 3d**) indicated the presence of at least two epitope clusters on the NTD, including one targeted by all of the tested IGHV1-24 mAbs (4A8, CM25, CM17, CM58, and 1-68) and the IGHV3-11 mAb CM30. Within this epitope cluster, we saw particularly strong levels of competition among CM17, CM25, CM58, and 4A8 (Fig.3d, dashed box). Another NTD epitope was identified by CM31 (IGHV2-5, 6.4% mutation), which overlaps with CM30 (IGHV3-11; 3.1% mutation), CM58, and 1-68 but does not compete with the other three IGHV1-24 NTD mAbs. Interestingly, CM31 instead showed low-level competition with the RBD-specific mAb CM32.

### Structural convergence of IGHV1-24 NTD antibodies

To better understand the IGHV1-24 NTD-reactive antibody interactions with the spike, we determined a cryo-EM structure of CM25 bound to S-ECD at an overall resolution of 3.3 Å (**Fig. 4a; Extended Data Figs. 7-8**). Three CM25 Fabs are bound to each trimeric S-ECD protein via interactions with the NTD, but the flexibility of the NTD relative to the rest of the spike resulted in poor density for the CM25 interface. Focused refinement of the NTD–CM25 region was performed to improve map quality, enabling building and analysis of the binding interface. The structure revealed a heavy-chain–dominant mode of binding, with substantial contacts mediated by interactions between the three CDRs and the N3 and N5 loops of the NTD (**Fig. 4b**). The light chain contributes 11% (86 Å^2^) of the total CM25 binding interface, mainly through a stacking interaction between CDR-L2 Tyr55 and Pro251 within the N5 loop. CDR-H1 interacts extensively through hydrogen bonds and hydrophobic interactions, including a salt bridge formed between the conserved Glu36 residue and the N5 loop residue Arg246 (**Fig. 4c**). As IGHV1-24 is highly electronegative (pI = 4.6), electrostatic interactions might be expected to guide the initial antibody–NTD association through steering and the formation of salt bridges and hydrogen bonds. The common IGHV1-24 Phe56 residue in CDR-H2 forms a pi-cation interaction with Lys147 in the N3 loop (**Fig. 4c**). CM25 contains a 14-amino-acid CDR-H3 loop that contributes 35% (261 Å^2^) of the total interface, including the AV aliphatic motif found in all but one of the convergent IGHV1-24 NTD-binding mAbs. Ala109 and Val110 are buried at the interface in a binding pocket framed by the N3 and N5 loops. The one extant structure of an IGHV1-24 NTD-binding antibody isolated by B-cell cloning was recently determined for mAb 4A8^3^. A comparison of CM25 with 4A8 reveals that the AV dipeptide interaction is structurally conserved, and the 21 amino-acid CDR-H3 of 4A8 extends along the outside of the NTD, contributing 3 additional contacts and 46% (415 Å^2^) of the total binding interface (**Fig. 4d**). Both structures show extensive contacts between the heavy chain of the Fabs and the N3 and N5 loops of the NTD. Notably, the Glu36-Arg246 salt bridge and an identical CDR-H2 contact between Phe56 and Lys147 are also observed at the 4A8-NTD interface.

**Figure 4.**
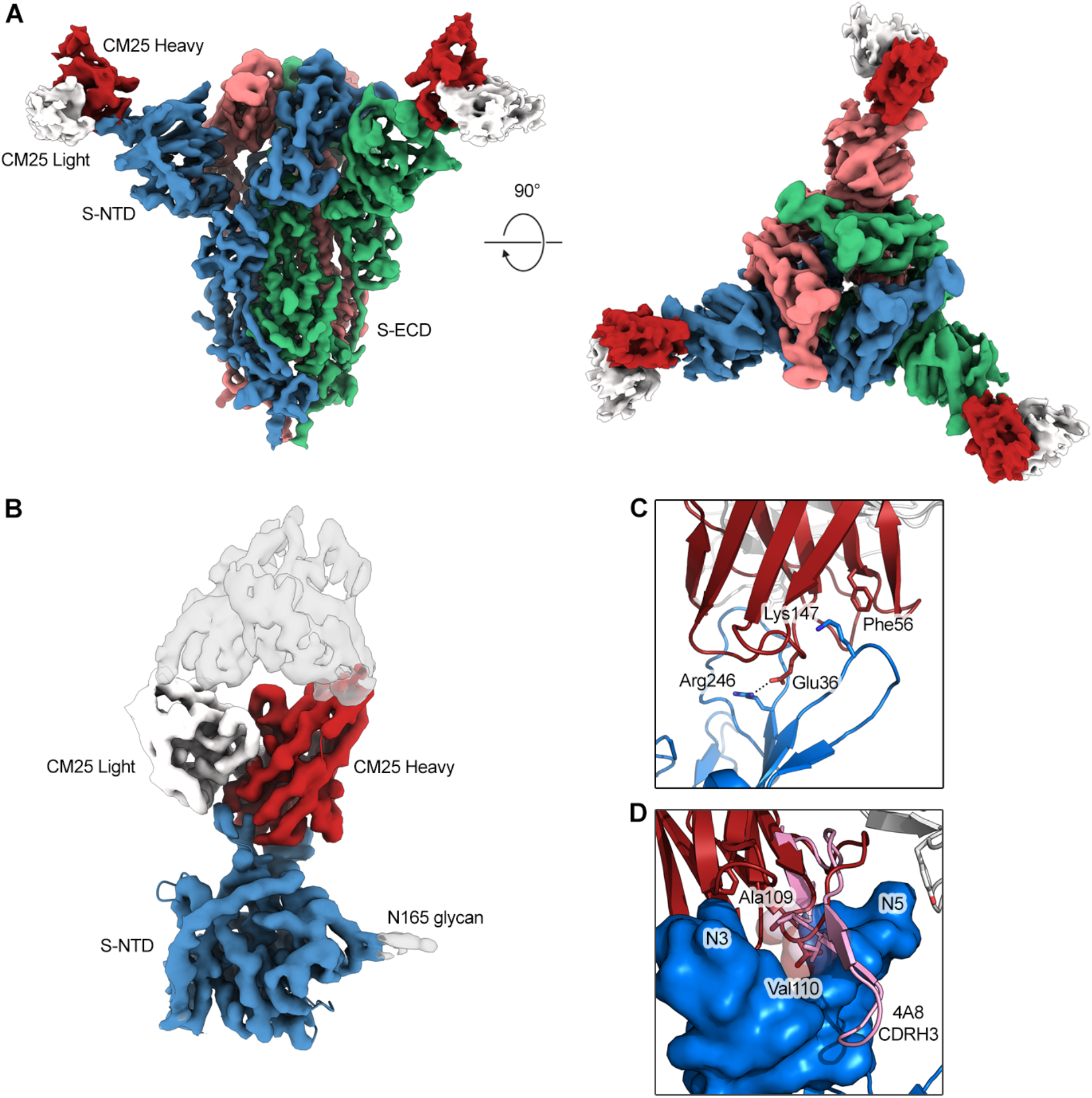
Structural basis of a shared, or public, class of IGHV1-24 plasma antibodies targeting the spike NTD. **a**, Side and top views of the structure of CM25 Fab bound to S-ECD shown as cryo-EM density. **b**, Focused refinement density revealing a VH-dominant mode of binding, with substantial contacts mediated by interactions between the three CDRs and the N3 and N5 loops of the NTD. **c**, CDR-H1 interaction includes a salt bridge formed between the uniquely encoded Glu36 residue and the N5 loop residue Arg246; Phe56 unique residue in CDR-H2 forms a pi-cation interaction with Lys147 in the N3 loop. **d**, The AV dipeptide interaction with the N3 and N5 loops of the NTD is structurally conserved between mAbs CM25 (red) and 4A8 (pink).

## DISCUSSION

Using molecular antibody proteomics to ascertain the S protein regions targeted by plasma antibodies in the response to SARS-CoV-2, we observe that the response is restricted in diversity (6–22 IgG lineages) and is directed predominantly to non-RBD spike epitopes in the four study subjects we have analyzed. This result appears to contradict a prior report^20^ claiming ∼90% of the SARS-CoV-2 neutralizing humoral immune response is accounted for by RBD-directed antibodies; however, nine of the 21 plasmas analyzed in that study nonetheless possessed 10%– 50% non-RBD-directed pseudovirus neutralizing activity; furthermore, the correlation of pseudovirus neutralizing activity with anti-RBD titers and ACE2-blocking activity was robust only for hospitalized patients with severe disease but, puzzlingly, not for non-hospitalized and presumably convalescent subjects (as we have studied here). Consistent with our results, serological profiling has identified a disconnect between plasma neutralizing titers and anti-RBD B-cell immunity in many individuals^21^. Also, a convalescent plasma was recently described^22^ wherein the dominant antibody (or antibodies) responsible for the majority of neutralizing activity target a single epitope in the NTD—which overlaps with the epitope bound by the common antibody class we have described here. To the best of our knowledge, our report is the first confirmation of the capacity of anti-NTD antibodies to confer protection *in vivo* and is the first demonstration of convergent antibody recognition of a spike epitope that resides outside the RBD, which collectively point to alternative routes for virus neutralization, the resolution of disease, and provide a rationale for therapeutic interventions based on non-RBD spike epitopes.

---

**Extended Data Table 1:**
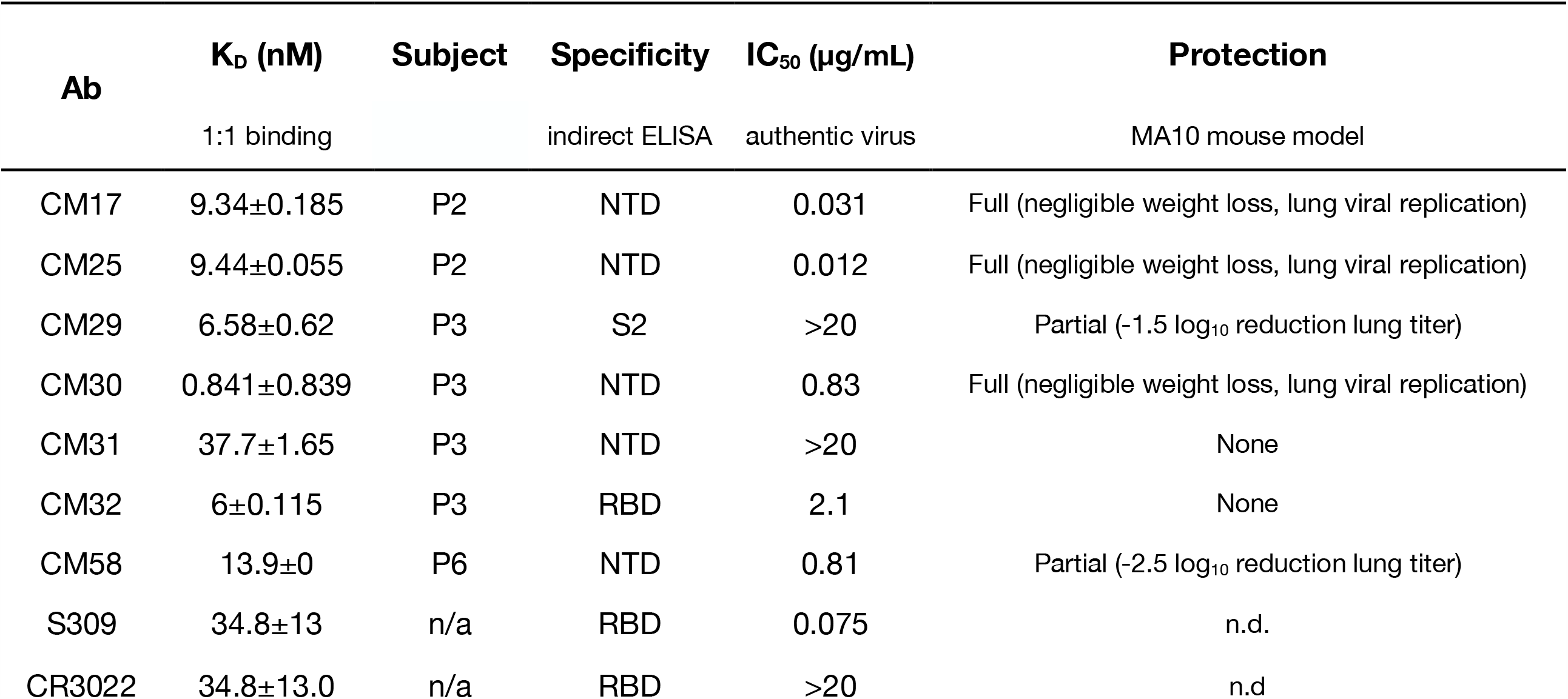
Recombinant plasma mAb binding and function.

| Ab | K<sub>d</sub> (nM) | Subject | Specificity | IC<sub>50</sub> (μg/mL) | Protection |
| --- | --- | --- | --- | --- | --- |
|  | 1:1 binding |  | indirect ELISA | authentic virus | MA10 mouse model |
| CM17 | 9.34±0.185 | P2 | NTD | 0.031 | Full (negligible weight loss, lung viral replication) |
| CM25 | 9.44±0.055 | P2 | NTD | 0.012 | Full (negligible weight loss, lung viral replication) |
| CM29 | 6.58±0.62 | P3 | S2 | >20 | Partial (-1.5 log<sub>10</sub> reduction lung titer) |
| CM30 | 0.841±0.839 | P3 | NTD | 0.83 | Full (negligible weight loss, lung viral replication) |
| CM31 | 37.7±1.65 | P3 | NTD | >20 | None |
| CM32 | 6±0.115 | P3 | RBD | 2.1 | None |
| CM58 | 13.9±0 | P6 | NTD | 0.81 | Partial (-2.5 log<sub>10</sub> reduction lung titer) |
| S309 | 34.8±13 | n/a | RBD | 0.075 | n.d. |
| CR3022 | 34.8±13.0 | n/a | RBD | >20 | n.d |

**Extended Data Table 2:**
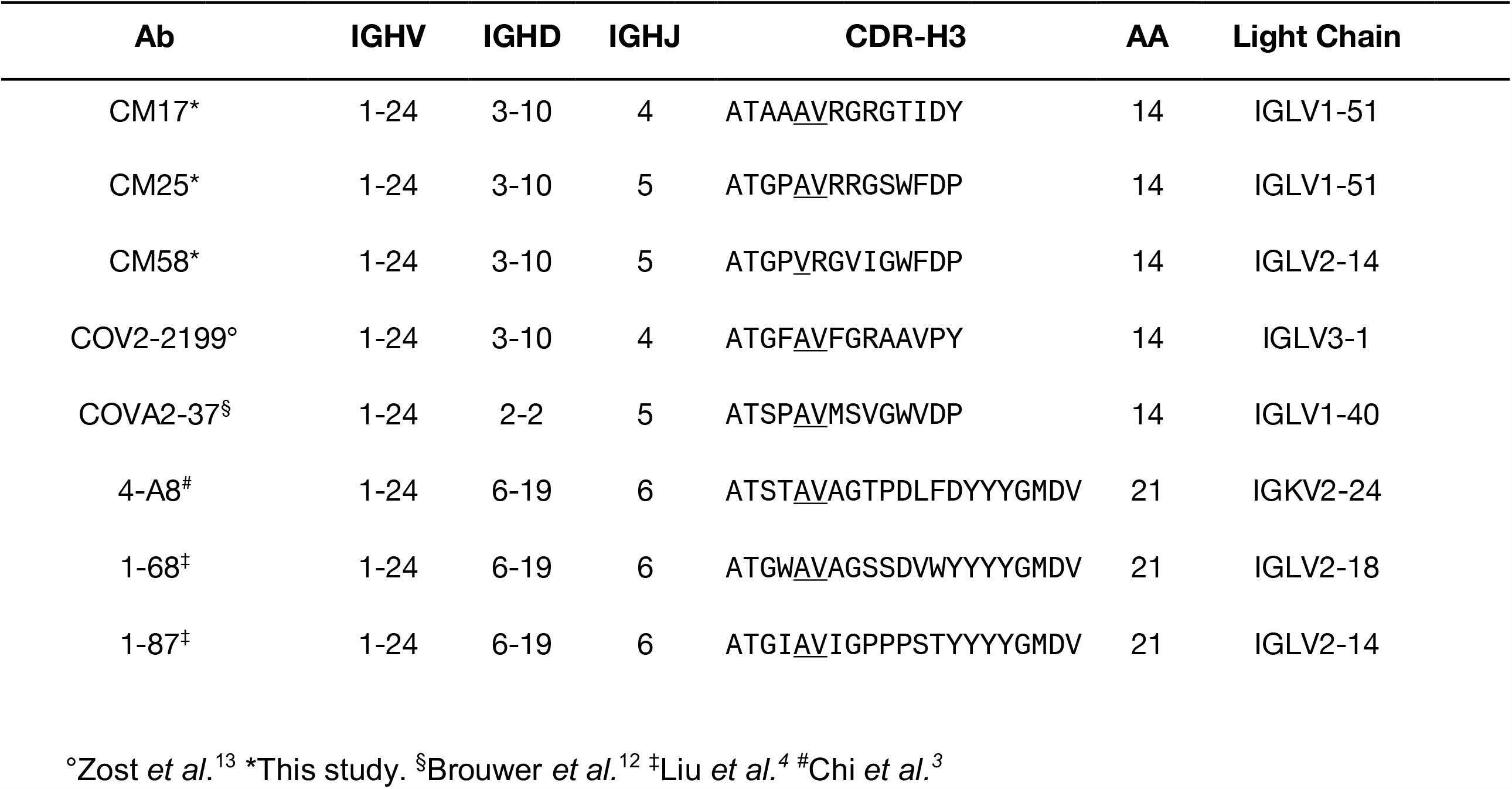
Convergent NTD mAbs.

**Extended Data Table 3:**
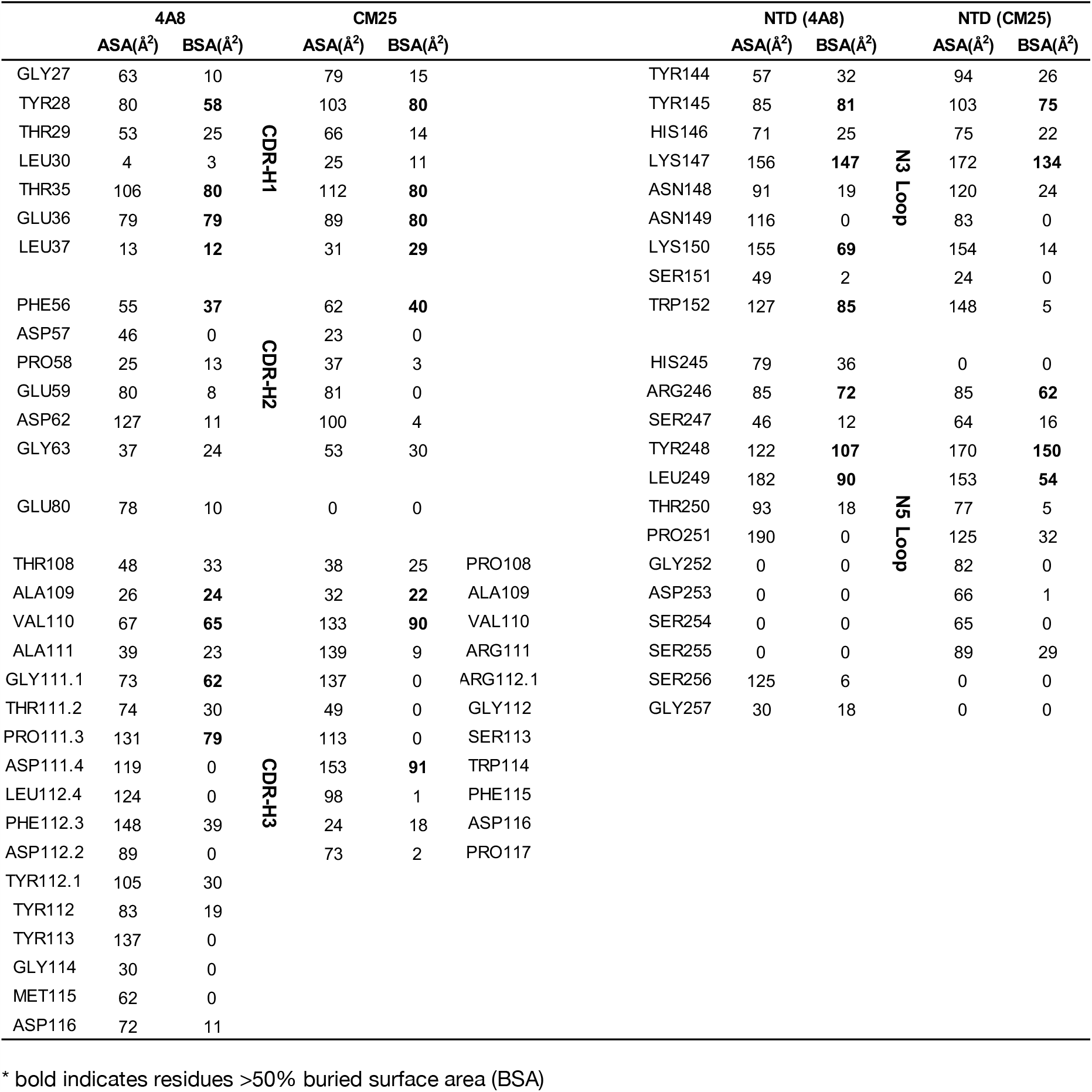
mAb–NTD interface analysis.

**Extended Data Table 4.**
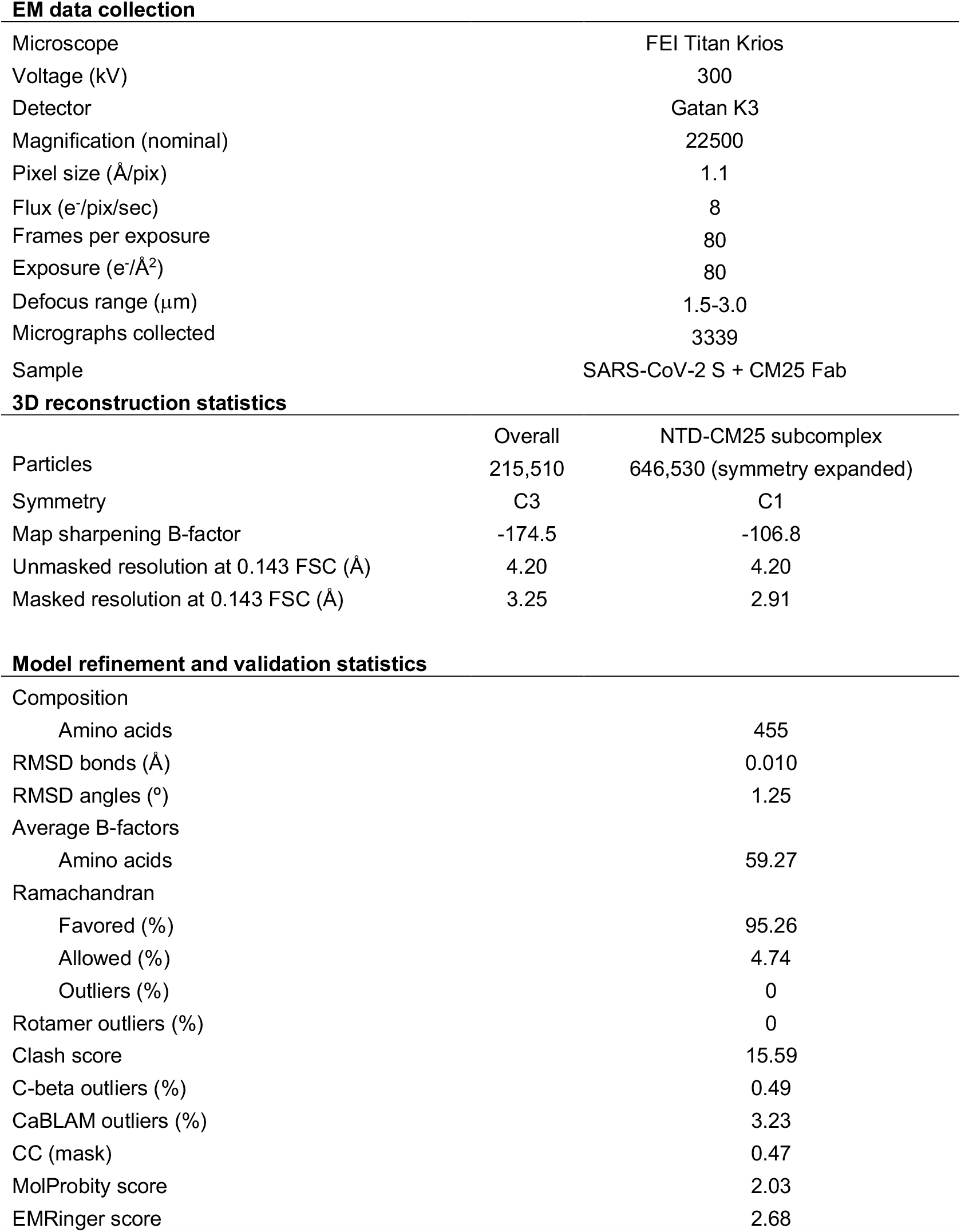

EM data collection
|  |  |
| --- | --- |
| Microscope | FEI Titan Krios |
| Voltage (kV) | 300 |
| Detector | Gatan K3 |
| Magnification (nominal) | 22500 |
| Pixel size (Å/pix) | 1.1 |
| Flux (e<sup>-</sup>/pix/sec) | 8 |
| Frames per exposure | 80 |
| Exposure (e<sup>-</sup>/Å<sup>2</sup>) | 80 |
| Defocus range (μm) | 1.5-3.0 |
| Micrographs collected | 3339 |
| Sample | SARS-CoV-2 S + CM25 Fab |

3D reconstruction statistics
|  | Overall | NTD-CM25 subcomplex |
| --- | --- | --- |
| Particles | 215,510 | 646,530 (symmetry expanded) |
| Symmetry | C3 | C1 |
| Map sharpening B-factor | -174.5 | -106.8 |
| Unmasked resolution at 0.143 FSC (Å) | 4.20 | 4.20 |
| Masked resolution at 0.143 FSC (Å) | 3.25 | 2.91 |
| Model refinement and validation statistics |  |
| Composition |  |
| Amino acids | 455 |
| RMSD bonds (Å) | 0.010 |
| RMSD angles (°) | 1.25 |
| Average B-factors |  |
| Amino acids | 59.27 |
| Ramachandran |  |
| Favored (%) | 95.26 |
| Allowed (%) | 4.74 |
| Outliers (%) | 0 |
| Rotamer outliers (%) | 0 |
| Clash score | 15.59 |
| C-beta outliers (%) | 0.49 |
| CaBLAM outliers (%) | 3.23 |
| CC (mask) | 0.47 |
| MolProbity score | 2.03 |
| EMRinger score | 2.68 |

## METHODS

### Austin cohort and collection of peripheral blood

All of the SARS-CoV-2 immune plasmas used for this study were collected from non-hospitalized PCR-confirmed individuals who presented with symptomatic disease. Whole blood was collected from convalescent COVID-19 subjects while they were quarantined at home. Study Subjects P1 and P2 blood draws occurred at days 12 and 56 post-onset of symptoms; Subject P3 at day 11; Subject P4 at days 19 and 45. Plasma and PBMCs were separated and collected by density gradient centrifugation using Histopaque-1077 media (Sigma-Aldrich).

### Expression and purification of SARS-CoV-2 proteins

For Ig-Seq antibody proteomics, the cloning, expression, and purification of the prefusion-stabilized spike ectodomain (S-ECD “S-2P”; GenBank: MN908947) encoding residues 1–1208 and containing two proline substitutions at 986 and 987 as well as other modifications, and residues 319–591 encoding the receptor binding domain (RBD), have been previously described^1^. For cryo-EM, spike protein was expressed by transiently transfecting plasmid encoding the HexaPro spike variant^2^ containing substitutions S383C and D985C^3^ with a C-terminal TwinStrep tag into FreeStyle 293-F cells (Thermo Fisher) using polyethyleneimine, with 5 μM kifunensine being added 3h post-transfection. The cell culture was harvested four days after transfection and the medium was separated from the cells by centrifugation. Supernatants were passed through a 0.22 µm filter followed by passage over StrepTactin resin (IBA). The sample was further purified by size-exclusion chromatography using a Superose 6 10/300 column (GE Healthcare) in buffer containing 2 mM Tris pH 8.0, 200 mM NaCl and 0.02% NaN_3_.

### ELISA

The methods for enzyme-linked immunosorbent assay to measure titers of anti-SARS-CoV-2 IgG plasma antibodies have been previously described^4^. For determination of mAb domain-level reactivity against recombinant spike ECD, RBD and NTD proteins, a standard indirect ELISA was used. Costar high binding 96-well assay plates (Corning) were coated with antigens (4 µg ml^-1^) in PBS. Antigens included in-house produced SARS-COV-2 spike ECD^2^ (S-ECD), SARS-COV-2 spike RBD, as well as commercially obtained SARS-COV-2 spike NTD (Sino Biological). Antigen-reactive mAbs were detected with goat anti-human IgG (Fab)-horseradish peroxidase (Sigma-Aldrich) conjugate in 1:5000 PBS. After washing with PBST-0.1%, the bound antibody was detected with 3,3′,5,5′-tetramethylbenzidine soluble substrate (TMB; Millipore) using a Synergy H1 Microplate Reader (BioTek Instruments, Inc.).

### V_H_ repertoire sequencing

PBMCs were lysed in TRIzol Reagent (Invitrogen) and total RNA was extracted using RNeasy (Qiagen). First strand cDNA was synthesized from 500 ng mRNA using SuperScript IV (Invitrogen), and cDNA encoding the V_H_ regions of the IgG, IgA, and IgM repertoires was amplified with a multiplex primer set^5^ using the FastStart High Fidelity PCR System (Roche) under the following conditions: 2 min at 95 °C; 4 cycles of 92 °C for 30 s, 50 °C for 30 s, 72 °C for 1 min; 4 cycles of 92 °C for 30 s, 55 °C for 30 s, 72 °C for 1 min; 22 cycles of 92 °C for 30 s, 63 °C for 30 s, 72 °C for 1 min; 72 °C for 7 min; hold at 4 °C, as previously described^5^. Products were sequenced by 2×300 paired-end Illumina MiSeq.

### Paired V_H_:V_L_ repertoire sequencing

PBMCs were co-emulsified with oligo d(T)25 magnetic beads (New England Biolabs) in lysis buffer (100mM Tris pH 7.5, 500mM LiCl, 10mM EDTA,1% lithium dodecyl sulfate, and 5mM dithiothreitol) using a custom flow-focusing device as previously described^6^. The magnetic beads were washed, resuspended in a one-step RT-PCR solution with an overlap extension V_H_ and V_L_ primer set as previously described^29^, emulsified using a dispersion tube (IKA), and subjected to overlap-extension RT-PCR under the following conditions: 30 min at 55 °C followed by 2 min at 94 °C; 4 cycles of 94 °C for 30 s, 50 °C for 30 s, 72 °C for 2 min; 4 cycles of 94 °C for 30 s, 55 °C for 30 s, 72 °C for 2 min; 32 cycles of 94 °C for 30 s, 60 °C for 30 s, 72 °C for 2 min; 72 °C for 7 min; hold at 4 °C. Amplicons were extracted from the emulsions, further amplified using a nested PCR, and sequenced using 2×300 paired-end Illumina MiSeq.

### Ig-Seq sample preparation and mass spectrometry

Total IgG was isolated from 1 mL plasma using Protein G Plus Agarose (Pierce Thermo Fisher Scientific) affinity chromatography and cleaved into F(ab’)_2_ fragments using IdeS. SARS-COV-2 Spike-specific F(ab’)_2_ was isolated by affinity chromatography using recombinant antigen (1 mg SARS-CoV-2 S-2P or RBD) coupled to 0.05 mg dry NHS-activated agarose resin (Thermo Fisher Scientific) as follows. F(ab’)_2_ (10 mg/mL in PBS) was rotated with antigen-conjugated affinity resin for 1 hour, loaded into 0.5 mL spin columns, washed 12X with 0.4 mL Dulbecco’s PBS, and eluted with 0.5 mL fractions of 1% formic acid. IgG-containing elution fractions were concentrated to dryness in a speed-vac, resuspended in ddH_2_O, combined, neutralized with 1 M Tris / 3 M NaOH, and prepared for liquid chromatography–tandem mass spectrometry (LC-MS/MS) as described previously^7^ with the modifications that (i) peptide separation using acetonitrile gradient was run for 120 min and (ii) data was collected on an Orbitrap Fusion (Thermo Fisher Scientific) operated at 120,000 resolution using HCD (higher-energy collisional dissociation) in topspeed mode with a 3s cycle time.

### Bioinformatic analysis

Raw Illumina MiSeq output sequences were trimmed according to sequence quality using Trimmomatic^8^ and annotated using MiXCR^9^. Sequences with ≥ 2 reads were clustered into clonal lineages defined by 90% CDRH3 amino acid identity using USEARCH^10^. LC-MS/MS search databases were prepared as previously described^7^, using custom Python scripts (available upon request). MS searches, and MS data analyses were performed as previously described^7^, adjusting the stringency of the elution XIC:flowthrough XIC filter to 2:1.

### Antibody expression and purification

Cognate V_H_ and V_L_ antibody sequences of interest were ordered as gBlocks (Integrated DNA Technologies) and cloned into a customized pcDNA 3.4 vector containing a human IgG1 Fc region. V_H_ and V_L_ plasmids were mixed at 1:2 ratio and were transfected into Expi293F cells (Thermo Fisher Scientific), which were cultured at 37 °C and 8% CO_2_ for 5 days, then neutralized and centrifuged at 1000 × g for 10 min. Antibodies was isolated from filtered supernatants using Protein G Plus Agarose (Pierce Thermo Fisher Scientific) affinity chromatography, washed with 20 column volumes of PBS, eluted with 100 mM glycine-HCl pH 2.5, and neutralized with 1 M Tris-HCl pH 8.0. The antibodies were buffer-exchanged into PBS and concentrated using 10,000 MWCO Vivaspin centrifugal spin columns (Sartorius).

### Binding affinity and checkerboard competition by biolayer interferometry

Bio-Layer interferometry (BLI) assays were performed using an 8-channel Octet RED96e instrument (ForteBio). Anti-hIgG Fc Capture (AHC) Biosensors (ForteBio 18-5060) were used and the assay was performed at 25 °C with shaking at 1,000 rpm. For IgG1 mAb K_D_ measurement, antibodies were diluted to 7.5 μg/mL and immobilized onto biosensors. The serial diluted HexaPro^2^ (200 nM to 12.5 nM) was associated for 3 min and dissociated for 5 min. The K_D_ were calculated using a 1:1 binding with drifting baseline model in BIAevaluation software. To determine NTD binding epitopes, BLI assays were performed as previously described. Briefly, the checkerboard experiment was performed with Anti-hIgG Fc Capture (AHC) Biosensors (ForteBio Inc., 18-5060) at 25 °C with shaking at 1,000 rpm. The first antibody was captured at 40 µg/ml for 10min and blocked with 50 µg/ml IgG isotype control for 5min. Antigen (NTD, 100µg/ml) was associated for 5min and 40 µg/ml of second antibody was associated for 5 min. The ForteBio Octet Data Analysis Software 9.0 was used for all analyses.

### SARS-CoV-2 Microneutralization Assay

#### USAMRIID

ATCC Vero E6 cells were seeded on 96-well plates 24-hours prior to infection. MAbs were normalized, 3-fold serially diluted, and incubated with a pre-titrated amount of SARS-CoV-2 virus (SARS-CoV-2/MT020880.1 isolate) at 37 °C for 1 hr. The virus-antibody inoculum was added to the Vero E6 monolayers and incubated for 24 hrs. Cells were then formalin fixed, permeabilized, and stained with a SARS-CoV nucleocapsid-specific antibody. After counterstaining, the monolayer was imaged under immunofluorescence software analyzed to quantify the presence of the detected antigen.

#### UNC

SARS-CoV-2 WA1 molecular clone (GenBank accession MT461669) and nLuc virus were generated previously^11^. ATCC Vero E6 cells were seeded at 20,000 cells per well on 96-well plates prior to infection. MAb samples were tested at starting concentrations of 30–0.1 μg/ml and were serially diluted three-fold up to eight dilution spots. Diluted mAbs were mixed with 87 PFU/well WT-nLuc virus, incubated at 37 °C with 5% CO_2_ for 1 hour, and then, following incubation, the growth medium was removed and virus–antibody mixtures were added to the cells in duplicate. Virus-only controls were included in each plate. After a 48h incubation at 37 °C with 5% CO_2_, cells were lysed, and luciferase activity was measured using Nano-Glo Luciferase Assay System (Promega) according to the manufacturer’s specifications. Neutralization titers were defined as the sample dilution at which a 50% reduction in relative light units (RLU) was observed relative to the average of the virus-only control wells.

### Evaluation of mAb prophylactic efficacy in the MA10 mouse model

For the inhibition of SARS-CoV-2 in the standard laboratory BALB/c mouse model, a pathogenic mouse ACE2-adapted SARS-CoV-2 variant, MA10, was constructed previously^11,12^. At 12h before infection, twelve-month-old female BALB/c mice (n=5/group) were injected intraperitoneally with 200μg/mouse of mAb or PBS. The mice were infected intranasally with a lethal dose (10^5^ or 10^4^ PFU) of the MA10 virus. Body weight of individual mice was measured daily, and all the mice were euthanized at day 4 post-infection by isoflurane overdose. The right caudal lung lobe was harvested and preserved in PBS at −80°C. Viral titers in the lung tissue were measured by plaque assay on Vero E6 cells.

### EM sample prep and data collection

Purified spike and Fab CM25 were combined at a final concentration of 0.2 mg/mL and 0.8 mg/mL respectively. Following a 30-minute incubation on ice, the complex was deposited on Au-300 1.2/1.3 grids that had been plasma cleaned for 4 minutes in a Solarus 950 plasma cleaner (Gatan) with a 4:1 ratio of O2/H2. The excess liquid was blotted for 3 seconds with a force of −4 using a Vitrobot Mark IV (Thermo Fisher) and plunge frozen into liquid ethane. 3,339 micrographs were collected from a single grid with the stage at a 30° tilt using a Titan Krios (Thermo Fisher) equipped with a K3 detector (Gatan). Movies were collected using SerialEM at 22,500X magnification with a corresponding calibrated pixel size of 1.1 Å^2^/ pixel. A full description of the data collection parameters can be found in **Extended Data Table 4**.

### Cryogenic electron microscopy (cryo-EM)

Motion correction was performed in Warp^13^. Micrographs were then imported into cryoSPARC v2.15.0 for CTF-estimation, particle picking, 2D classification, *ab initio* reconstruction, heterogenous 3D refinement and homogenous refinement^14^. For the CM25–NTD interface, the data were subjected to focused refinement using C3 symmetry expansion and particle subtraction. The focused mask contained the NTD and CM25 Fab. To improve map quality, the focused refinement volumes were processed using the DeepEMhancer tool via COSMIC^2^ science gateway^15^. An initial model was generated by docking PDBID: 6VSB^1^ and a Fab model based on the CM25 sequence built using SAbPred ABodyBuilder^16^ into map density via UCSF Chimera^17^. The model was built further and iteratively refined using a combination of Coot, Phenix, and ISOLDE^18-20^. The full cryo-EM processing workflow and structure validation can be found in **Extended Data Figs. 7-8**.

### Statistics

GraphPad Prism version 9.0.0 (GraphPad Software Inc., La Jolla, CA, USA) was used to conduct statistical analyses. Non-parametric Mann–Whitney U test and analysis of variance on ranks (Kruskal–Wallis H test) were used to determine the statistical significance of population means between two or more groups, respectively. Statistical differences in MA10 mouse modeling were tested using a one-way ANOVA with Dunnet’s multiple comparisons test, comparing every group with the mock-challenge lung titers.

## Ethics statement

The acquisition of blood specimens from convalescent individuals was approved by the University of Texas at Austin Institutional Review Board (protocol 2020-03-085; Breadth of serum antibody immune responses prior to, or following, patient recovery in asymptomatic and non-severe COVID-19). Informed consent was obtained from all participants.

## Data availability

The sequences all CM IgG plasma mAbs have been deposited in GenBank (https://www.ncbi.nlm.nih.gov/genbank/) with accession numbers *<u>XXY–XXZ</u>*. Coordinates for the

CM25 antibody Fab in complex with trimeric spike ectodomain have been deposited to the Protein Data Bank as PDB *<u>#αβγδ</u>*. These structural data are presented in Fig. 4, Extended Data Table 3, and Extended Data Figs. 7-8.

## Author Contributions

Conceptualization: WNV, GG, JJL and GCI; Methodology: WNV, YJH, NVJ, JEK, GD, APH, BLI, MDP, JD, AH, RSB, JSM, GG, JJL and GCI; Investigation: WNV, YJH, NVJ, JEK, GD, APH, FB, CP, YT, SAA, WP, KG, DRB, DMT, JG, DB, MG, JJL and GCI; Data Analysis and Interpretation: WNV, YJH, NVJ, JEK, GD, APH, SAA, WP, DRB, JRM, LR, DB, JL, JP, SG, SS, AH, JDG, RSB, JSM, GG, JJL and GCI; Data Curation: WNV, JYH, NVJ, JEK, GD, JJL and GCI; Writing: Original Draft, WNV, NJV, JJL, and GCI; Writing: Review & Editing: WNV, YJH, NJV, RSB, JSM, GG, JJL and GCI; Funding: JDG, RSB, JSM, GG and GCI.

## Acknowledgements

We are indebted to our study subjects for providing the blood samples required for this study. We wish to thank Dr. Gregory Fenves, former President of The University of Texas at Austin, Dr. Daniel Jaffee, Vice President for Research, and Dr. Paul Goldbart, Dean of the College of Natural Sciences, for their support. The authors are grateful for the tireless administrative expertise of Elizabeth K. Miller, to The LaMontagne Center for Infectious Disease, and for the university’s superb core facilities during this trying period. Funding for USAMRIID was provided through the CARES Act with programmatic oversight from the Military Infectious Diseases Research Program–project 14066041. Opinions, conclusions, interpretations, and recommendations are those of the authors and are not necessarily endorsed by the U.S. Army. The mention of trade names or commercial products does not constitute endorsement or recommendation for use by the Department of the Army or the Department of Defense. The findings and conclusions in this report are those of the authors and do not necessarily represent the views of Centers for Disease Control and Prevention. Molecular graphics and analyses performed with UCSF Chimera, developed by the Resource for Biocomputing, Visualization, and Informatics at the University of California, San Francisco, with support from NIH P41-GM103311. The Sauer Structural Biology Laboratory is supported by the University of Texas College of Natural Sciences and by award RR160023 from the Cancer Prevention and Research Institute of Texas (CPRIT). This research was funded in part by the Clayton Foundation (G.G.); a National Institutes of Health (NIH)/National Institute of Allergy and Infectious Diseases (NIAID) grant awarded to J.S.M. (R01-AI127521); NIH NCI COVID-19 SeroNet grant U54-CA260543 (R.S.B.); and in part with federal funds under a contract from the NIH NIAID, Contract Number 75N93019C00050 (G.G., J.J.L., G.C.I.).

## EXTENDED DATA

**Extended Data Figure 1.**
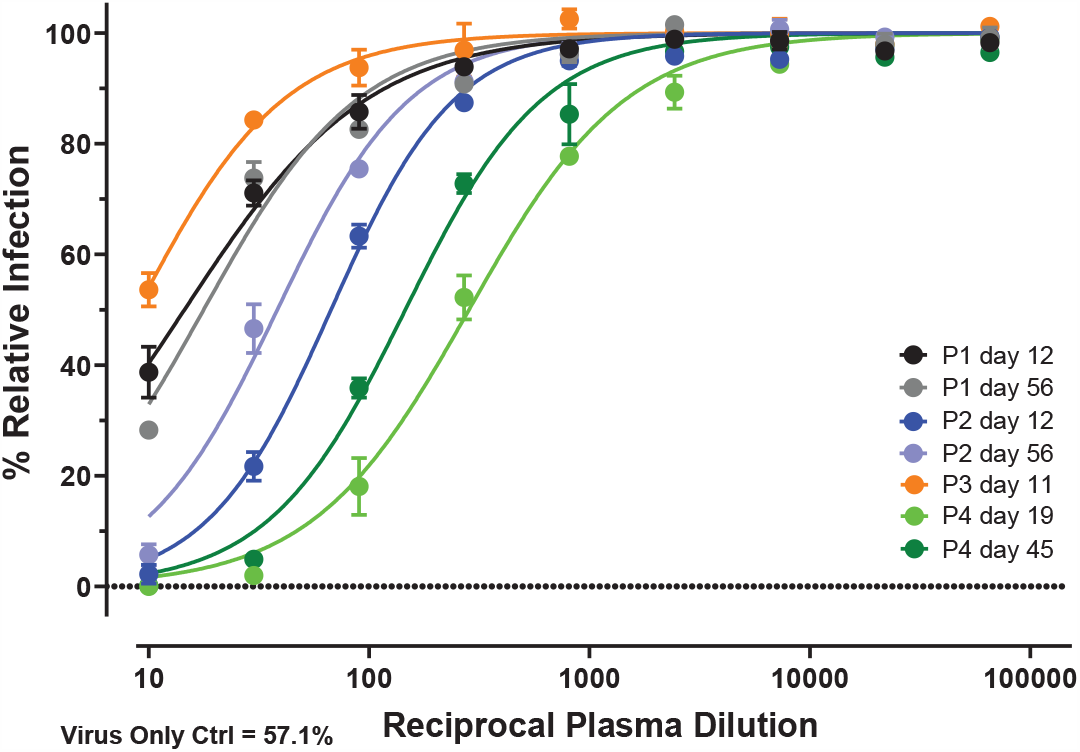
Live virus neutralization titers of four COVID+ study subjects’ plasma at each collection time point. Serial dilutions of plasma were tested in duplicate (SD error bars) for inhibition of live SARS-CoV-2 virus infection of in vitro monolayered Vero E6 cells. The percent of infected Vero E6 cells in each sample dilution was normalized relative to the virus-only (no plasma) negative control sample.

**Extended Data Figure 2.**
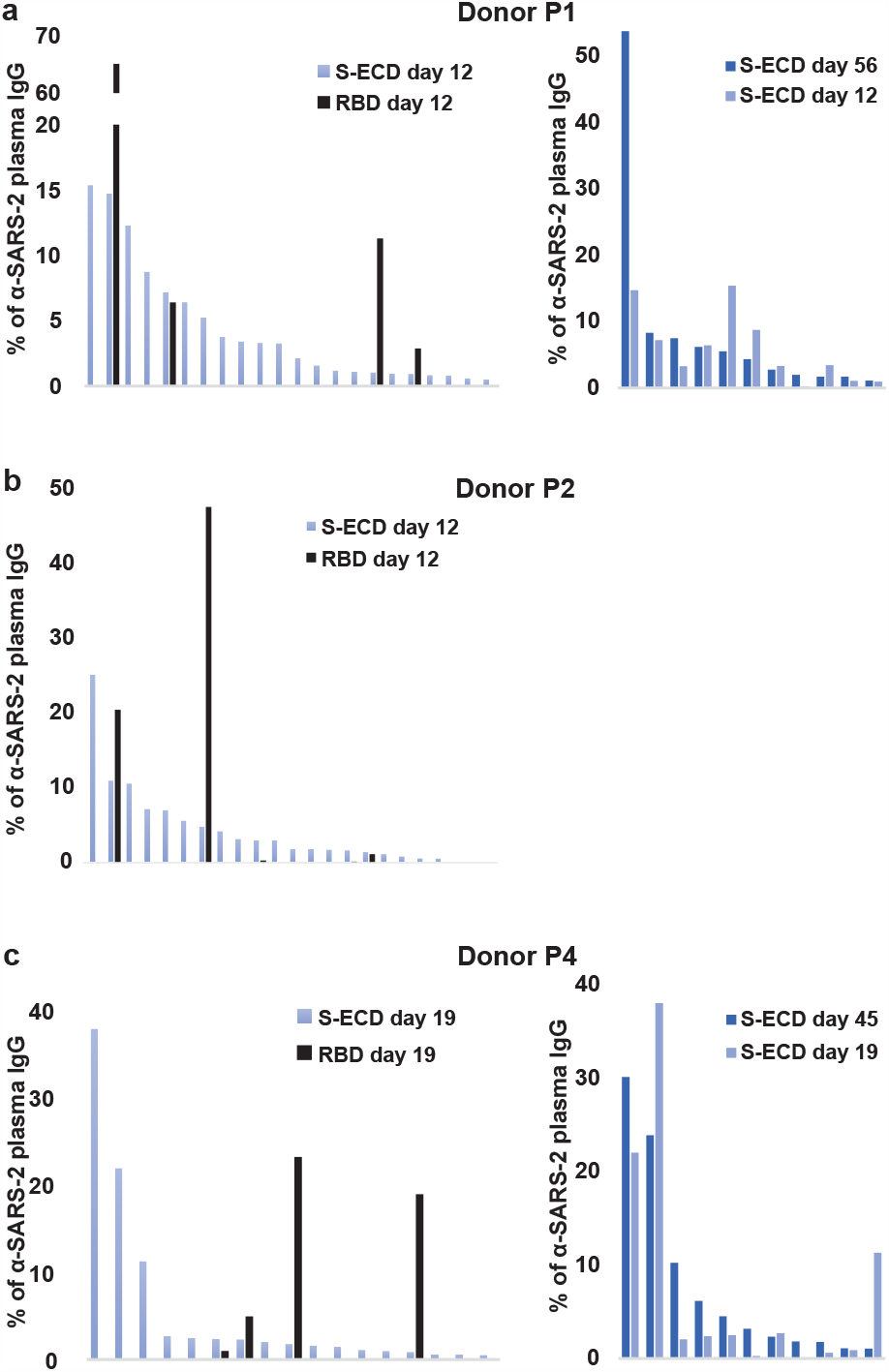
Ig-seq plasma IgG lineage profiles of study subjects at early and late convalescent time points. On the left, the first time point Ig-seq profile (days 11-19) for each subject (subject P3 found in Fig.1) shows both the SARS-CoV-2 spike ECD (S-ECD) and RBD abundance for each plasma IgG lineage detected at >0.5% anti-S-ECD plasma IgG (summed lineage XIC). Similarly, on the right, the second time point data for S-ECD (days 45-56) is shown for each lineage detected at >0.5% S-ECD plasma IgG abundance (time point 2), alongside early time point S-ECD data for comparison (subject P2 found in Fig.2).

**Extended Data Figure 3.**
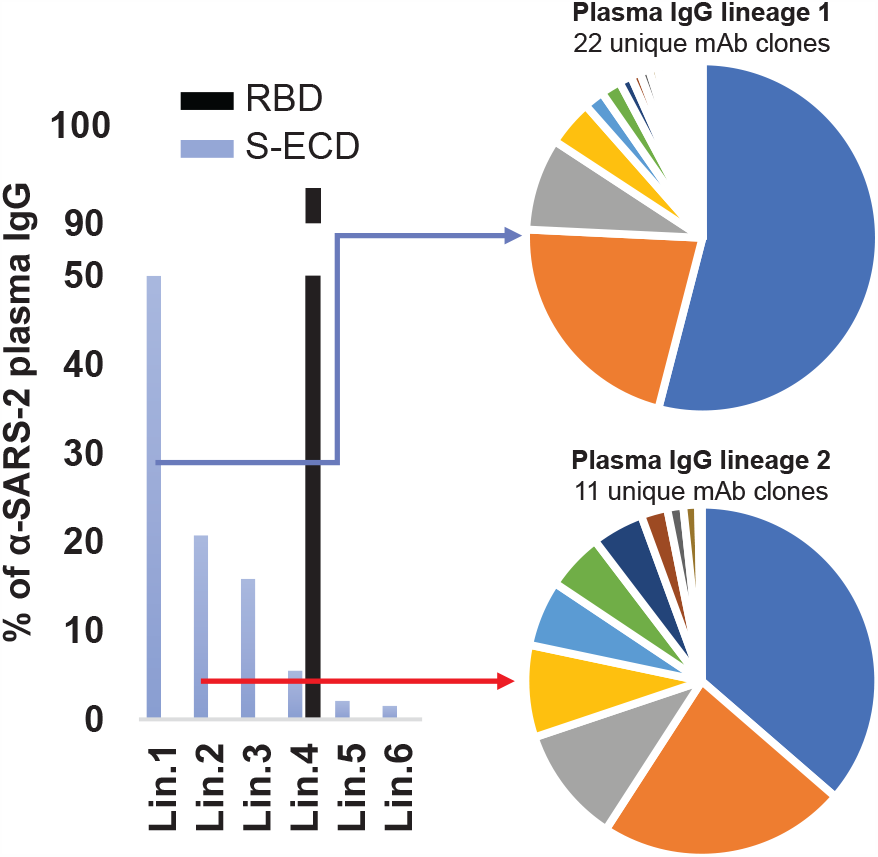
Ig-seq intra-lineage diversification in study subject P3 at day 11. The top two plasma IgG lineages from subject P3 demonstrate a large number of LC-MS/MS identified unique CDR-H3 clones within each lineage (33 total unique CDR-H3 clones in top two IgG lineages combined). This indicates extensive ongoing diversification within this donor at early convalescence.

**Extended Data Figure 4.**
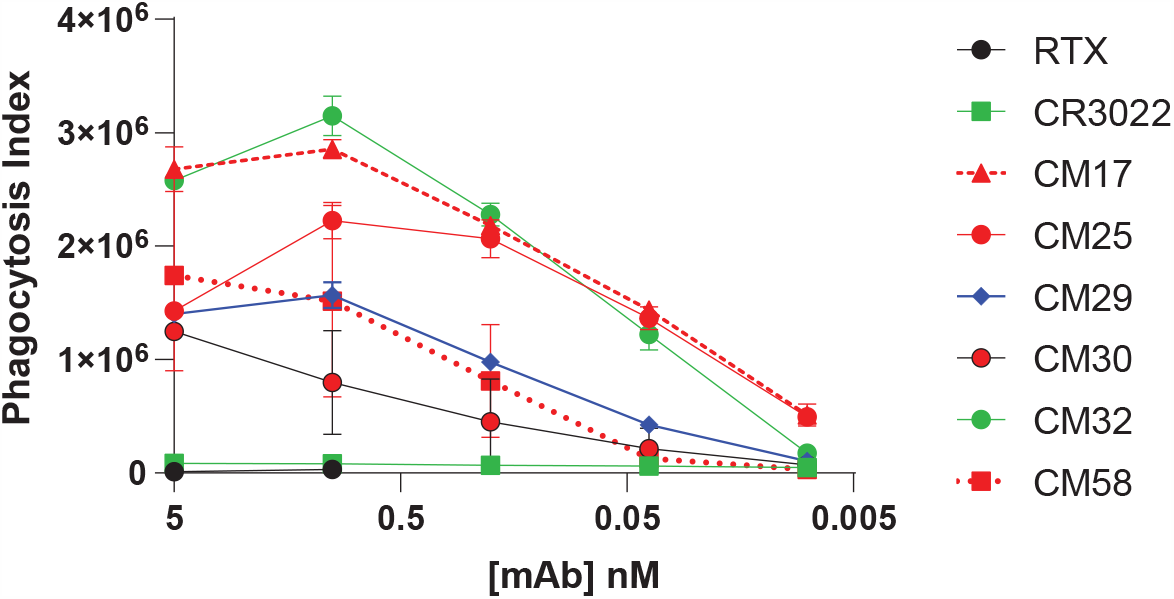
ADCP ECD-bead assay using recombinant plasma IgG mAbs. ADCP activity of recombinant plasma mAbs serially diluted on THP-1 cells in the presence of spike-conjugated, fluorescent polystyrene beads. The phagocytosis index metric represents the percent of bead-positive THP-1 cells multiplied by the average MFI of each cell to account for increased levels of bead internalization.

**Extended Data Figure 5.**
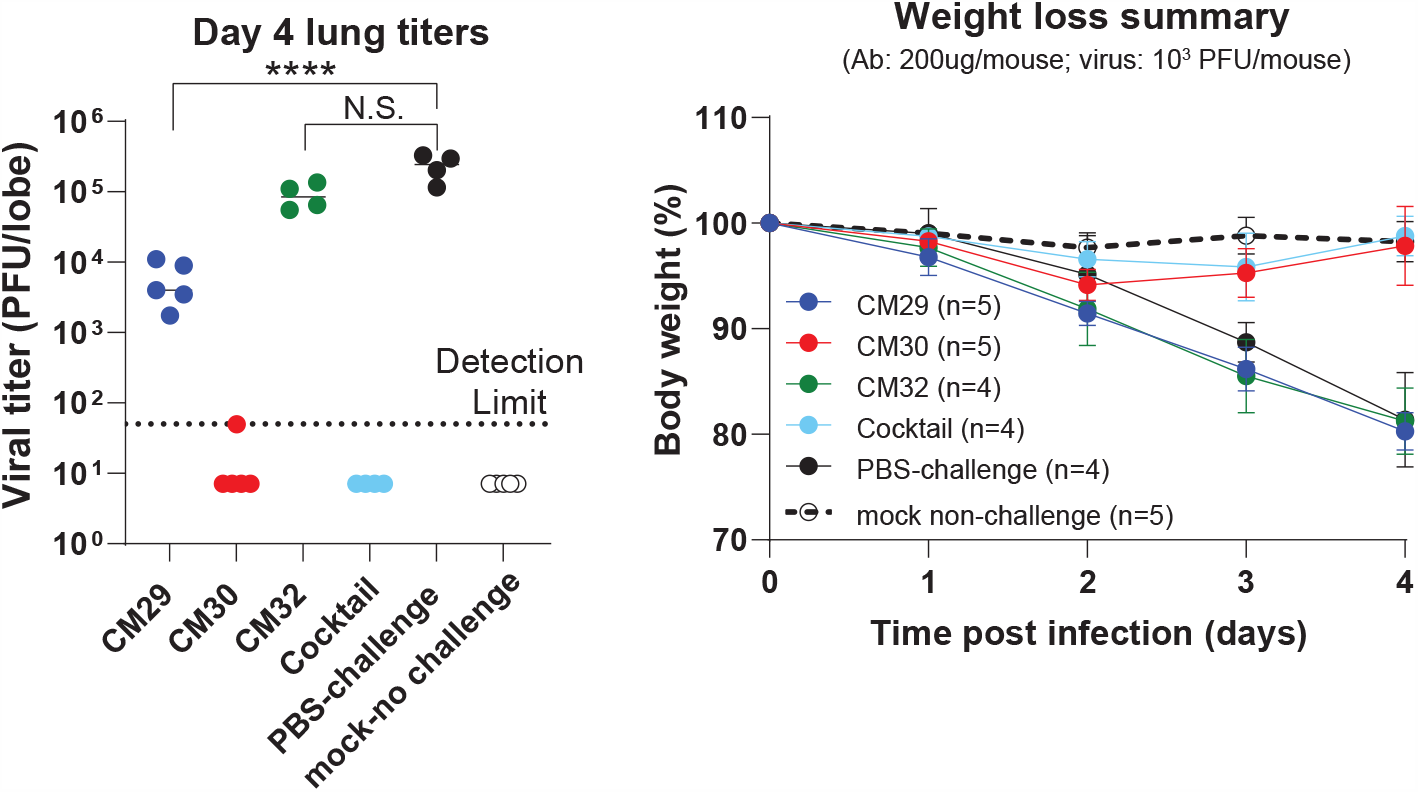
In vivo protection against SARS-CoV-2 viral challenge using recombinant plasma IgG mAbs. Day four lung viral titers and average cohort weight loss of 12 m.o. BALB/C mice after intranasal challenge with 10^3^ PFU of mouse-adapted (MA10) SARS-CoV-2. 200 µg of mAb was administered 12 hours prior to challenge. Cocktail consisted of all four of the top donor P3 plasma IgG lineages (CM29, CM30, CM31, CM32) in combination.

**Extended Data Figure 6.**
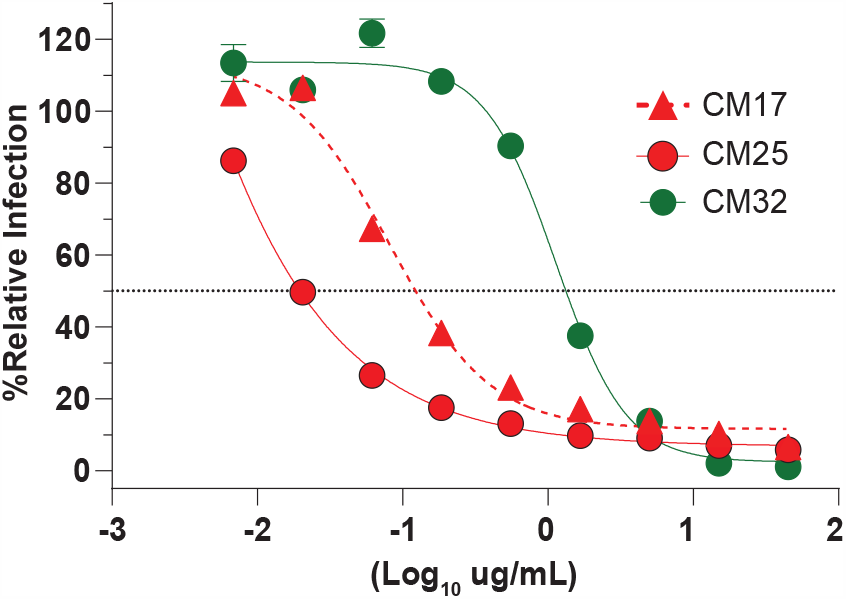
Independent live virus neutralization titers of recombinant plasma IgG mAbs CM17, CM25, and CM32. In vitro live virus neutralization curves for CM17, CM25, and CM32 repeated in second independent laboratory demonstrate similar levels of inhibition (as compared to data in Fig.1e and 2c) of live SARS-CoV-2 virus infection of monolayered Vero E6 cells. The percent of infected Vero E6 cells in each sample dilution was normalized relative to the virus-only (no plasma) negative control sample.

**Extended Data Figure 7.**
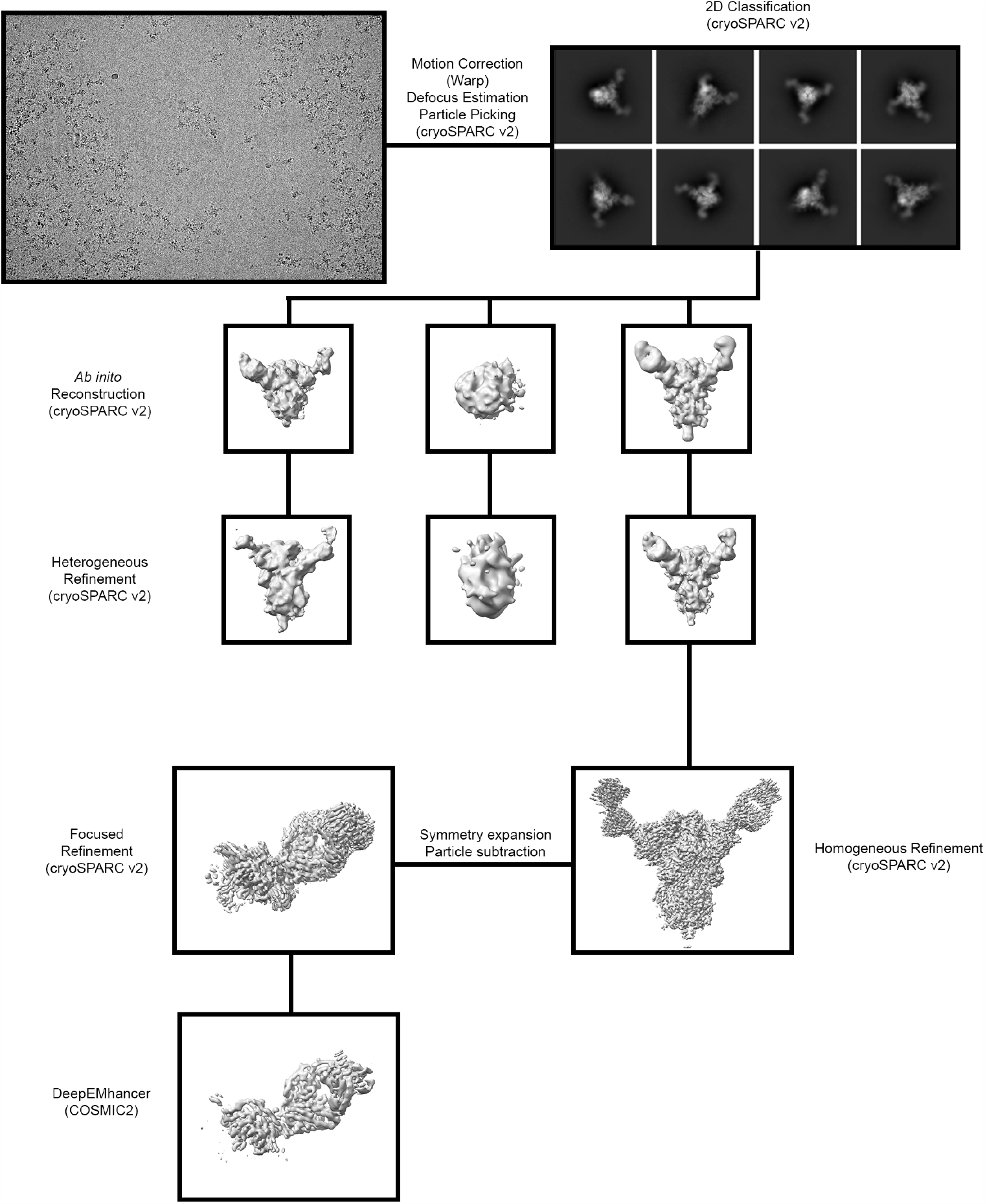
Cryo-EM data processing workflow.

**Extended Data Figure 8.**
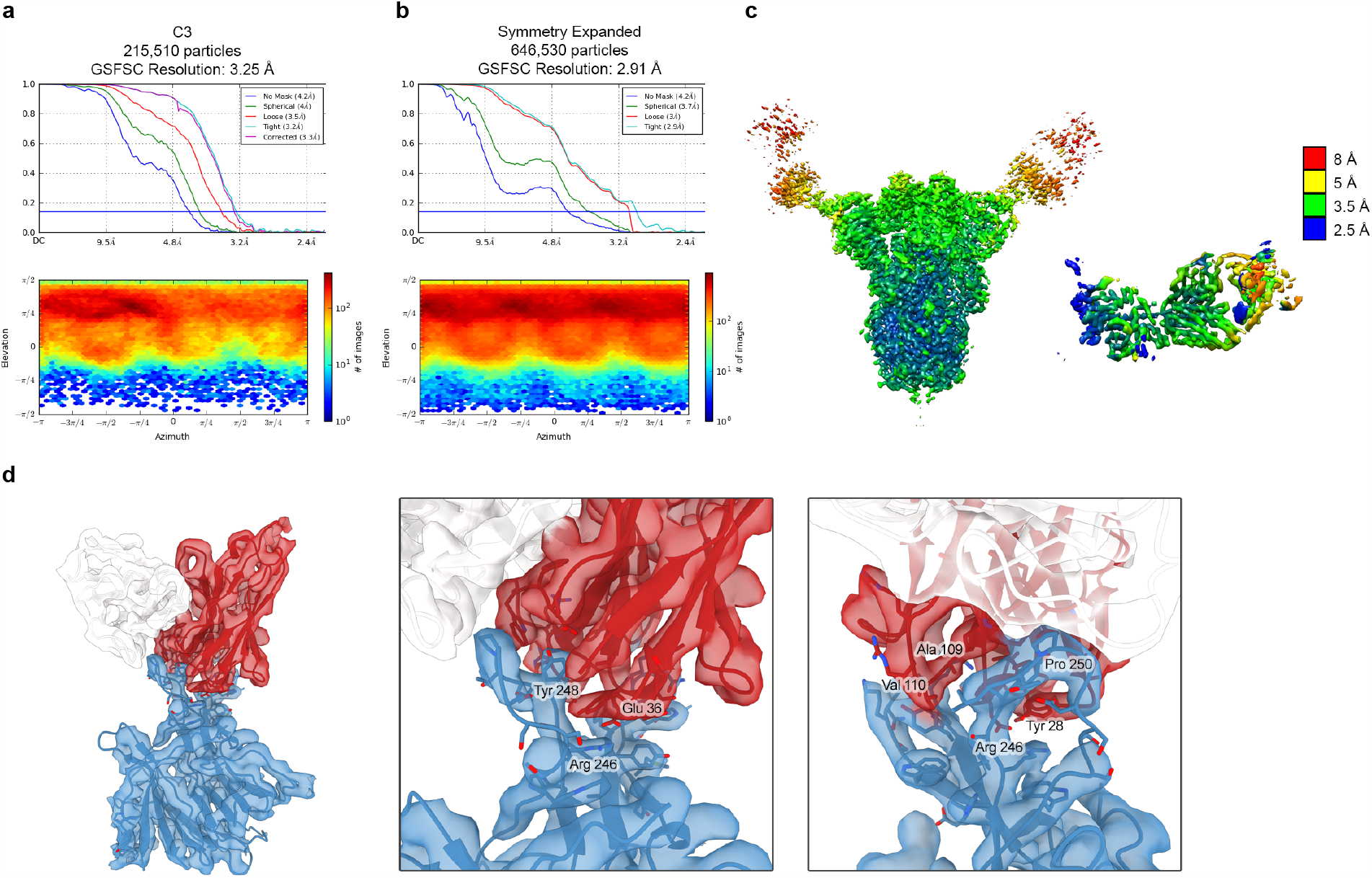
Cryo-EM structure validation. **a**, FSC curve and viewing distribution plot for the overall S-CM25 Fab structure, generated in cryoSPARC v2.15. **b**, FSC curve and viewing distribution plot for the focused refinement of S-NTD bound to the CM25 Fab. **c**, Cryo-EM density of the overall S-CM25 (left) and the focused S-NTD-CM25 (right) reconstructions, colored according to local resolution. **d**, Focused refinement density and corresponding models for the NTD (blue), CM25 Heavy chain (red), and CM25 Light chain (white). Full view of the NTD-CM25 Fab complex (left). Detailed views of the binding interface (middle, right). Oxygen atoms are colored red, nitrogen blue, and sulfurs yellow.

